# Enhanced Herd-Wide Surveillance Testing Strategy for Porcine Reproductive and Respiratory Syndrome Virus

**DOI:** 10.64898/2026.01.26.701801

**Authors:** Bailey M. Barcal, Joshua L. DeMers, Alison C. Neujahr, Christine E. Mainquist-Whigham, Jamie M. Madigan, Kristen K. Bernhard

**Affiliations:** BMB, JLD, ACN, KKB: DARO, Inc, Lincoln NE 68508; CEMW, JMM: Pillen Family Farms, Columbus NE

**Keywords:** swine, Porcine Reproductive and Respiratory Syndrome Virus, surveillance testing, whole-herd, oral fluid

## Abstract

**Objective:** This study aimed to compare a novel surveillance methodology to detect Porcine Reproductive and Respiratory Syndrome virus against oral fluid methodology in swine herds.

**Materials and methods:** Two pilot studies were conducted using two separate, high-risk commercial nurseries in central Nebraska, comparing two different surveillance sampling approaches (DARO Systems vs. oral fluid (OF)) in the detection of Porcine Reproductive and Respiratory Virus (PRRSV). Each nursery contained eight rooms with an average site inventory of 12,500 pigs. Weekly testing conducted in three of the eight rooms using DARO Systems and OF methodology to identify PRRSV until there was a positive sample, then daily testing of all rooms was conducted. Reverse Transcription-Quantitative Polymerase Chain Reaction was used for identification of positive PRRSV.

**Results:** Surveillance testing using novel methodology DARO Systems identified PRRSV in nurseries on average 3.91 days earlier than OF.

**Implications:** DARO Systems allows for a more robust whole-herd sampling technique to rapidly and accurately detect PRRSV 3.91 days earlier than gold standard approaches. Additionally, DARO Systems allows for an unbiased, whole-herd sampling approach. This method enables producers to implement earlier disease mitigation strategies.

---

Porcine reproductive and respiratory syndrome virus (PRRSV) is a highly infectious virus that causes one of the most disruptive diseases in the global swine industry in regards to biosecurity and animal welfare of infected animals.^1, 2^ PRRSV has been estimated to cost the US swine industry USD 1.2 billion/year due to the reduction of animal performance, higher mortality rates, and an increase in input cost for treatment and prevention.^1–3^ The need for accurate detection in surveillance testing is crucial for prompt treatment. Traditionally, blood collection from individual animals has been the most common sampling method.^1^ However, such sampling is tedious, and can be dangerous for the samplers as well as stressful for the pigs.^1^ Oral fluid collection has become the sampling method of choice due to its non-invasive approach and cost effectiveness.^1, 4, 5^ Oral fluids allow for the pigs to chew on an absorptive device such as a cotton rope or swab where the fluid is composed of saliva produced by the salivary glands and transudate.^1, 4, 6–9^ As a result, this fluid is composed of the presence of pathogens derived locally or from the circulatory system.^4^ Samples obtained from oral fluids can be collected at both the individual or group level and pooled for downstream testing. While oral fluids allows for a non-invasive sampling method, accurate data is dependent on infected animals chewing on the absorptive device and has shown to have variable success rate in accurate detection^10^. Since the early 2000’s, the implementation of oral fluid surveillance testing has allowed for swine producers to routinely test for problematic swine pathogens using a non-invasive methodology and ultimately has changed the way producers monitor disease in swine populations.^11^ As such, reports have shown the increasing trend of such sampling over the years, specifically testing Porcine Reproductive and Respiratory Syndrome virus (PRRSV).^11^ Oral fluid testing among the three major diagnostic labs within the US has seen an increase in PRRSV testing over a six-year span from roughly 14,000 test/annually to roughly 148,000 test/annually.^11^ However, producers are encouraged to exercise caution when using oral fluids as literature consistently shows variability performance with oral fluid matrix.^4, 11, 12^

Successful infectious disease surveillance programs require fast, convenient, and simple sampling procedures and laboratory testing. In addition, surveillance methodologies need to be accurate, cost-effective, animal welfare friendly, and user friendly.^1^ Thus, easy-to-use tools are needed to monitor and eliminate the widespread distribution and economic impact PRRSV has on the swine industry.^1^ Here we report a non-invasive, unbiased, convenient, and simple population surveillance testing procedure, known as the DARO Systems, that accurately detects PRRSV on average 3.91 days compared to traditional Gold Standard methods, such as oral fluid.

## Materials and methods

### Animals and Care

This study involved non-invasive environmental fecal sampling and analysis of retrospective oral fluid data collected as part of routine herd monitoring. No live animal handling or animal sampling was performed specifically for research purposes. Accordingly, review and approval by the DARO Institutional Animal Care and Use Committee (IACUC) were determined to be not required.

### Study Design

The study was conducted on a commercial swine farm using two high-risk commercial nurseries (Pilot 1 and Pilot 2) in central Nebraska. These two locations were selected to evaluate time of detection between two sampling methods in the face of a natural PRRSV infection within a site. Within both pilots, each nursery (Pilot 1 nursery barn and Pilot 2 nursery barn) contained eight rooms consisting of 24 pens stocked at 3 sq. feet. The estimated average site inventory was 12,500 pigs. Pigs were placed at 21 days of age and remained in the nursery for 7 weeks until transported to a finisher. Each of the rooms within both pilots was sampled weekly using both collection approaches. Specifically, one DARO Systems sample was taken from each room of the nursery during both pilots. Similarly, a pooled oral fluid sample was obtained per room for analysis. Upon a positive PRRSV test, daily samples were collected using both approaches until each produced a positive test until the time of shipment,

Pigs were PRRSV naïve at weaning with no intentional live virus exposure or modified live PRRSV vaccination administered. Based on area pressure, these barns were expected to be at high risk for PRRSV exposure during their turn. Rooms that tested positive for PRRSV by oral fluids and DARO Systems collection on day 1 of sampling were omitted (Table 3 & Table 4).

**Table 1.**
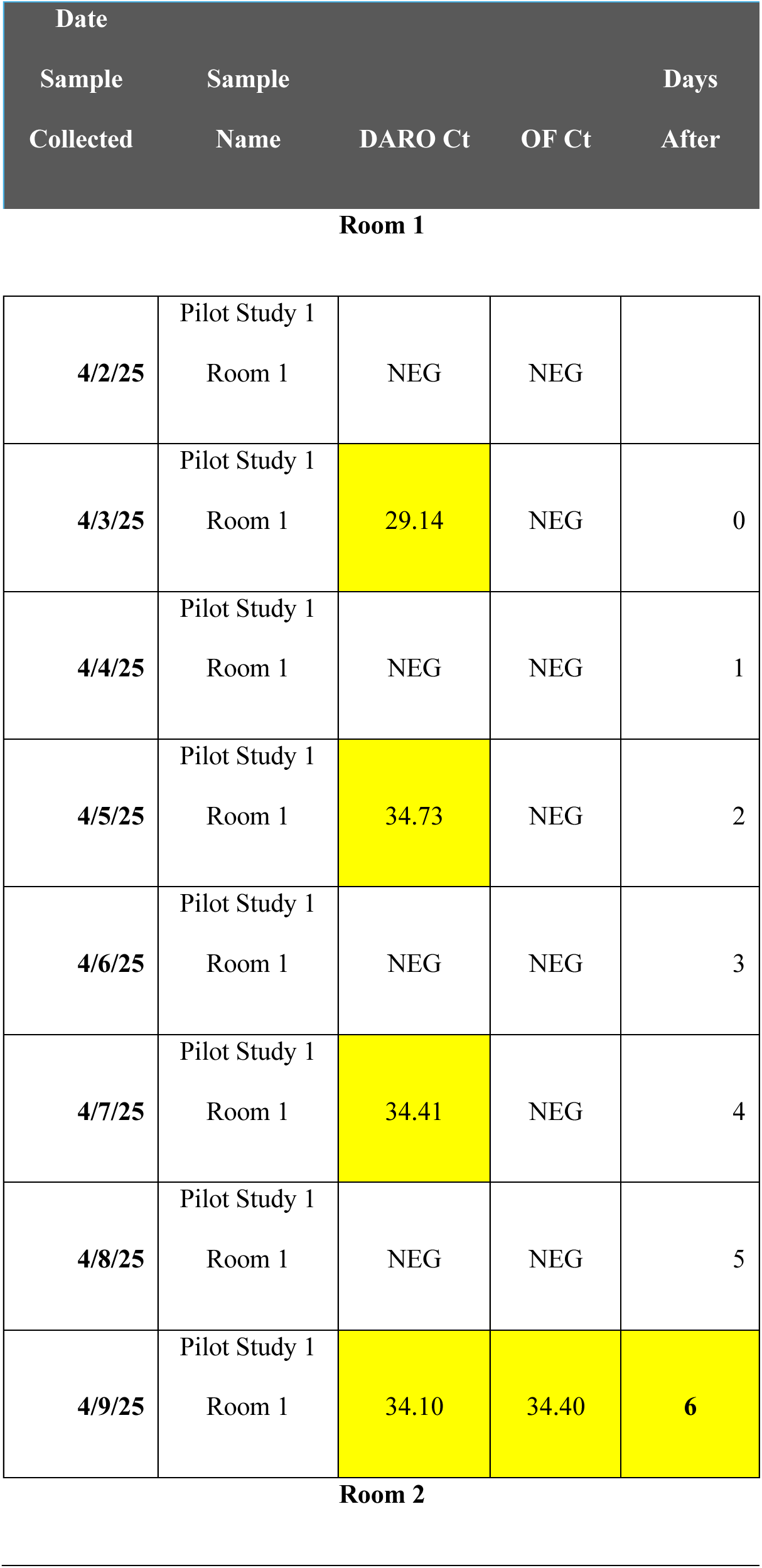
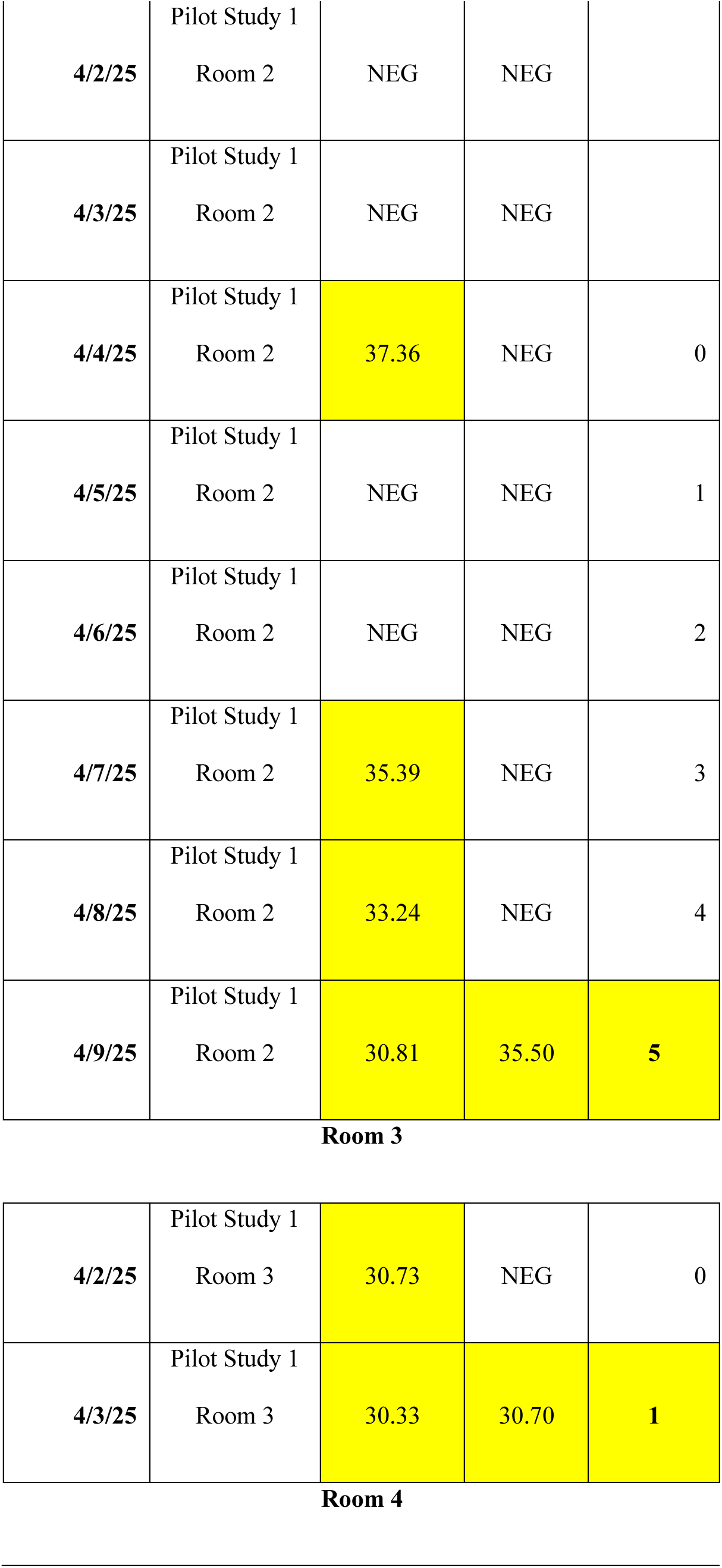
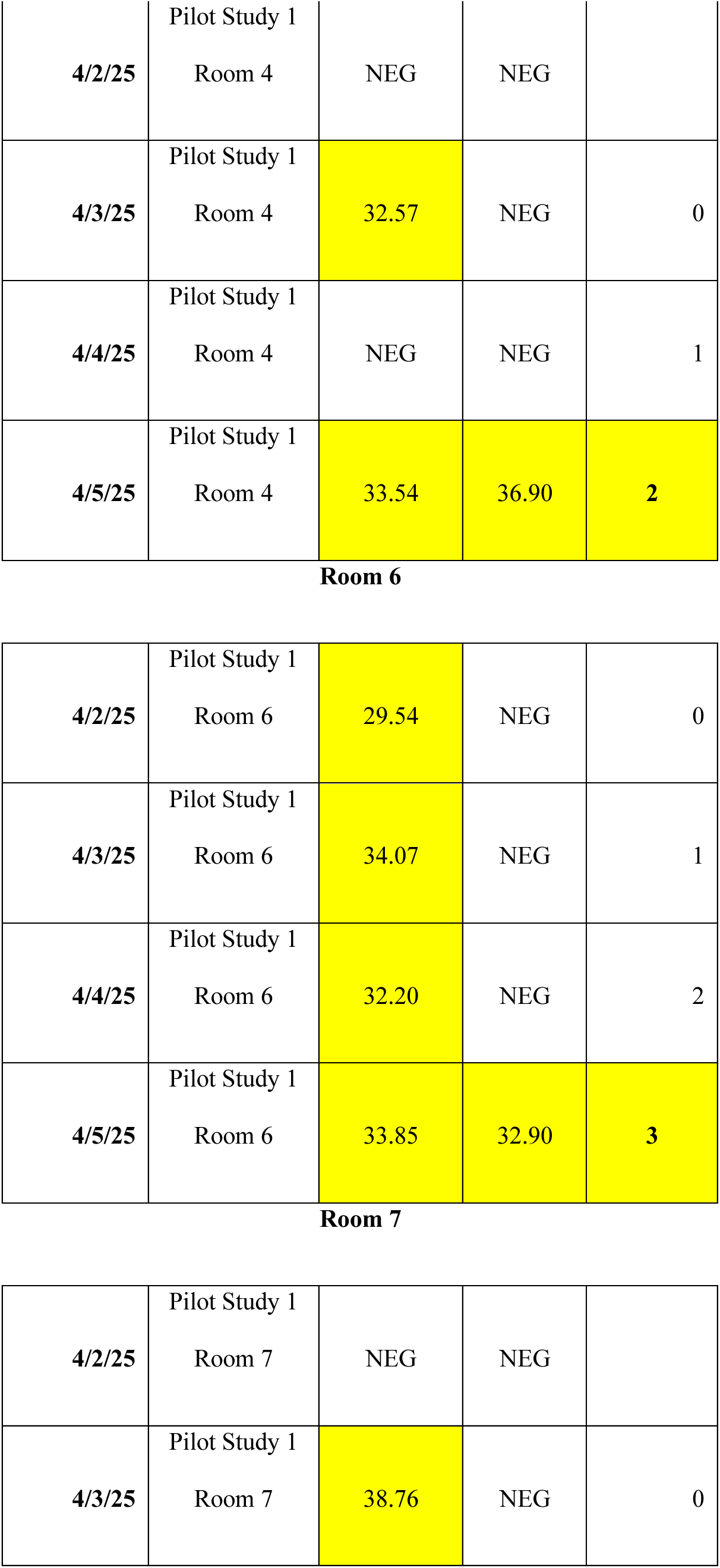
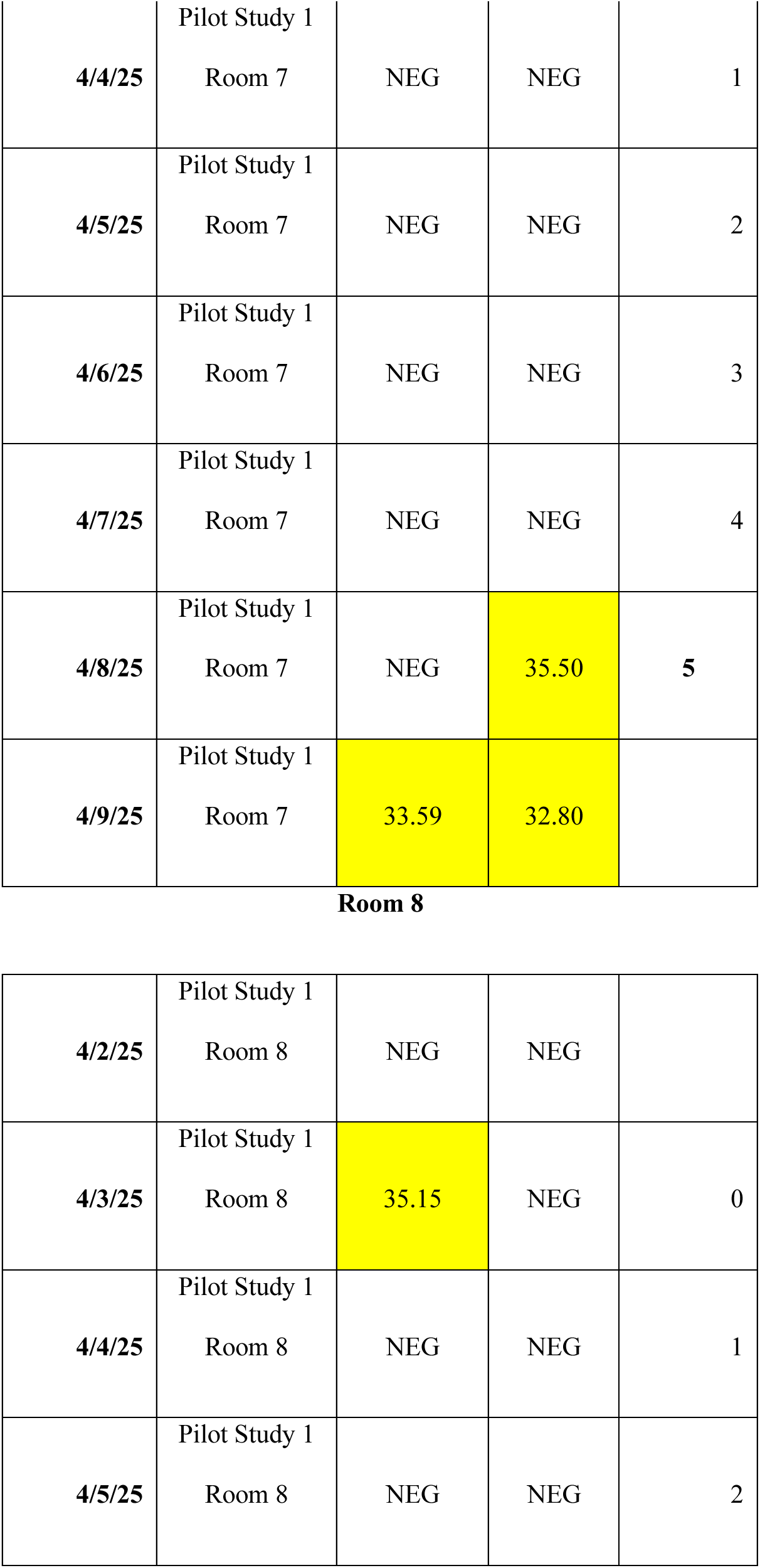
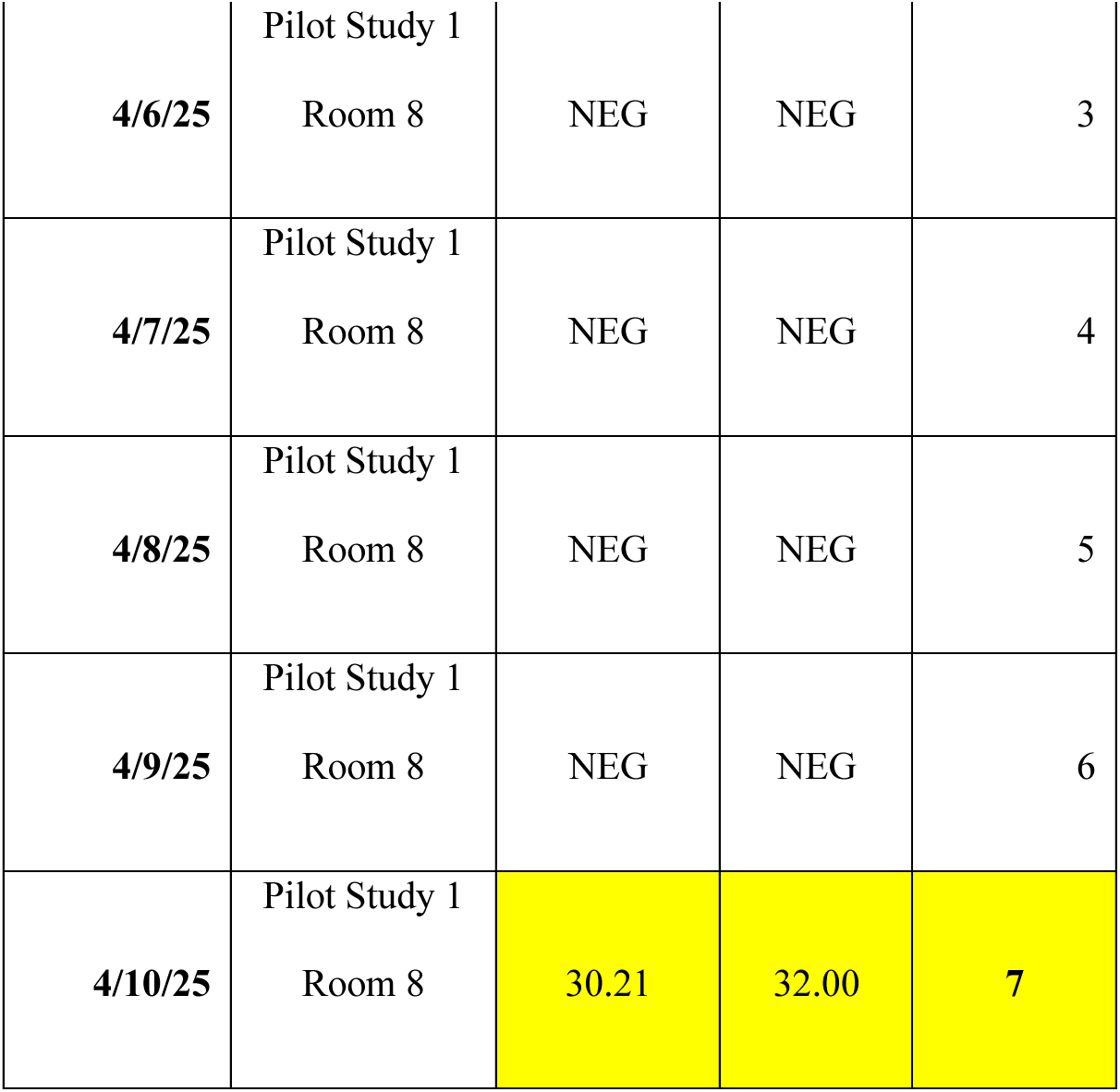
RT-qPCR Ct value report between oral fluid and DARO Systems testing among each room within Pilot Study 1. All data is shown until both surveillance methods Ct are deemed positive. Upon positive for both methodologies, sampling ceased.

**Table 2.**
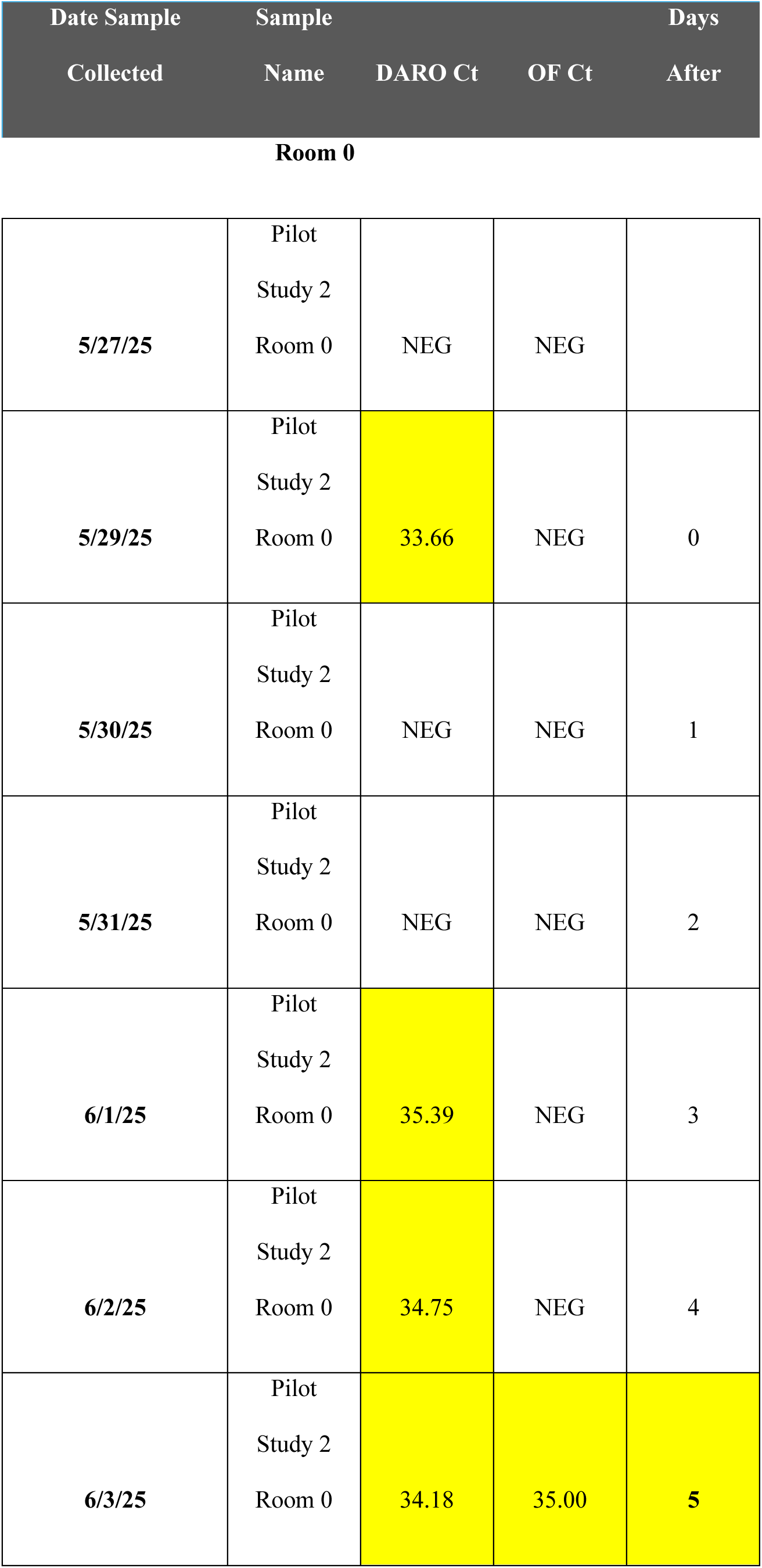
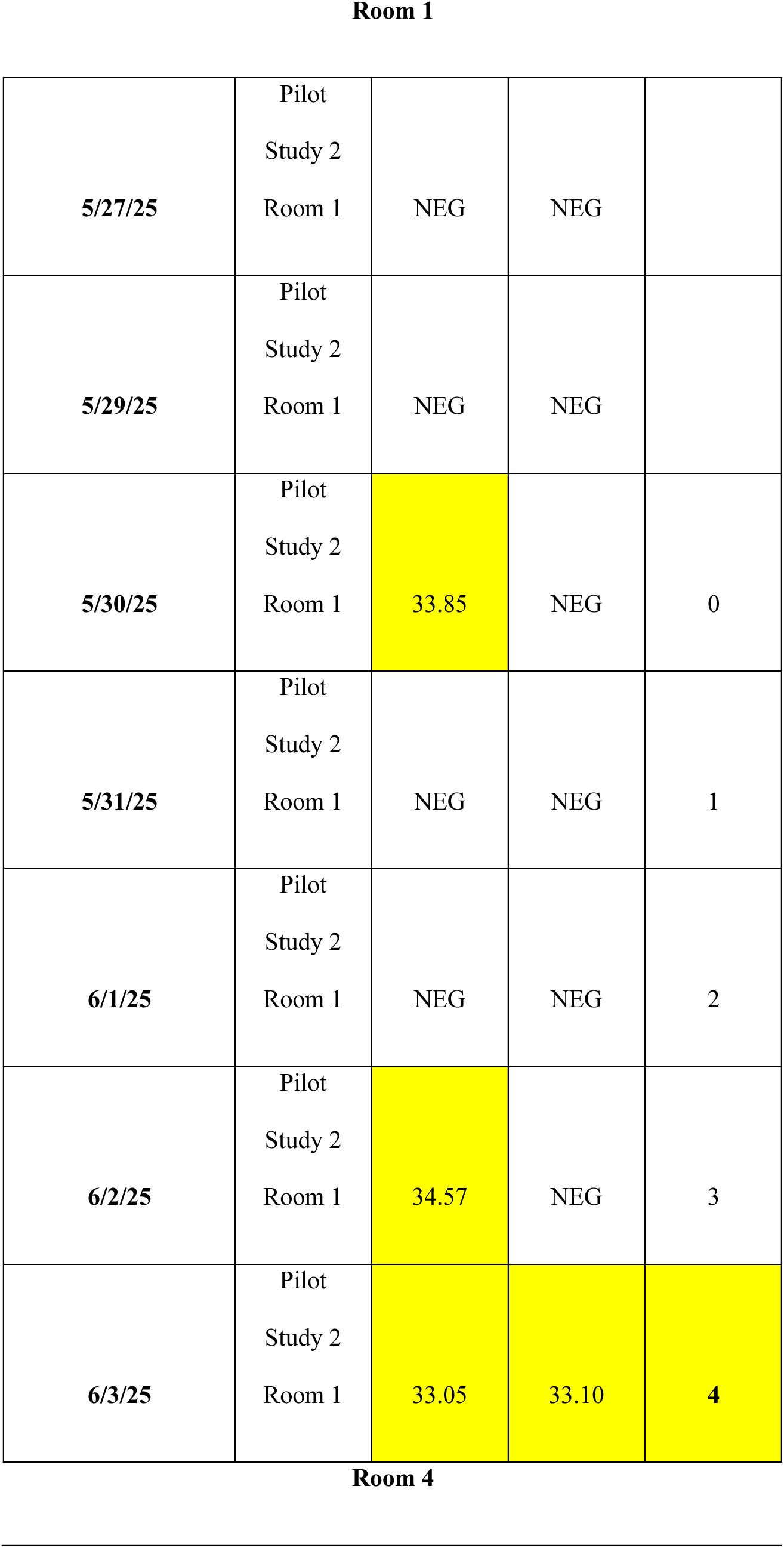
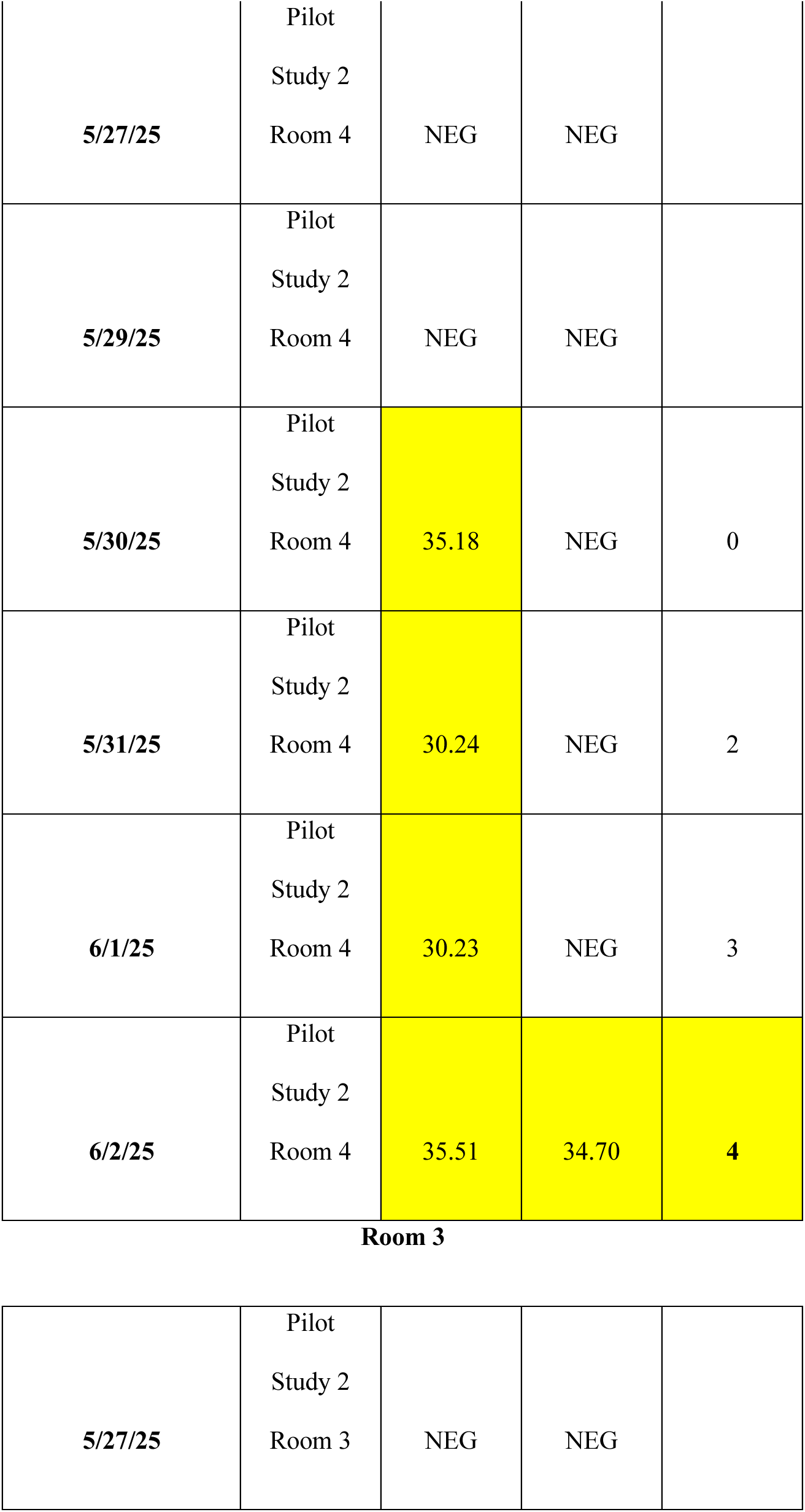
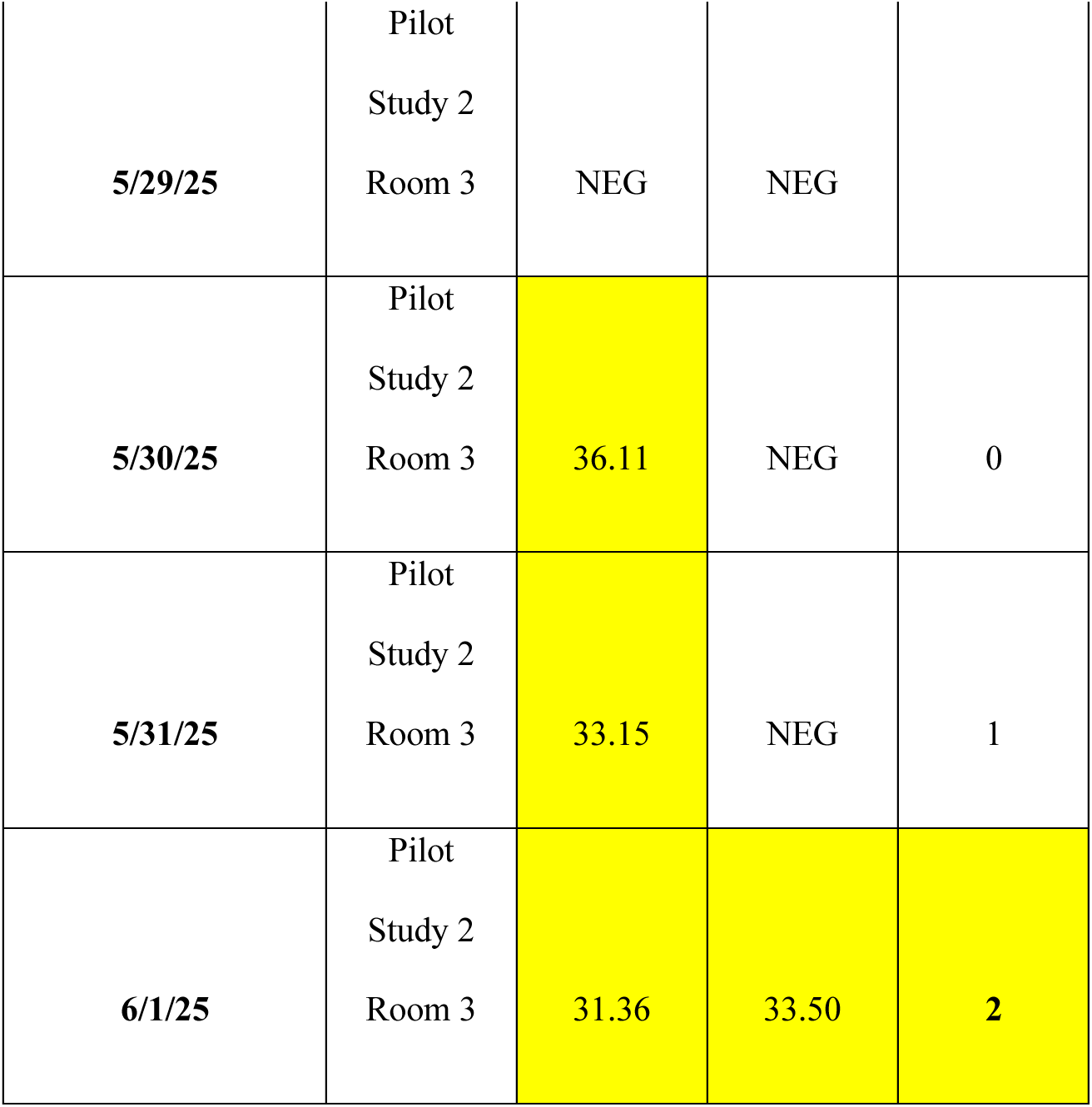
RT-qPCR Ct value comparison between oral fluid and DARO Systems testing among each room within Pilot Study 2. All data is shown until both surveillance methods Ct are deemed positive. Upon positive for both methodologies, sampling ceased.

**Table 3.**
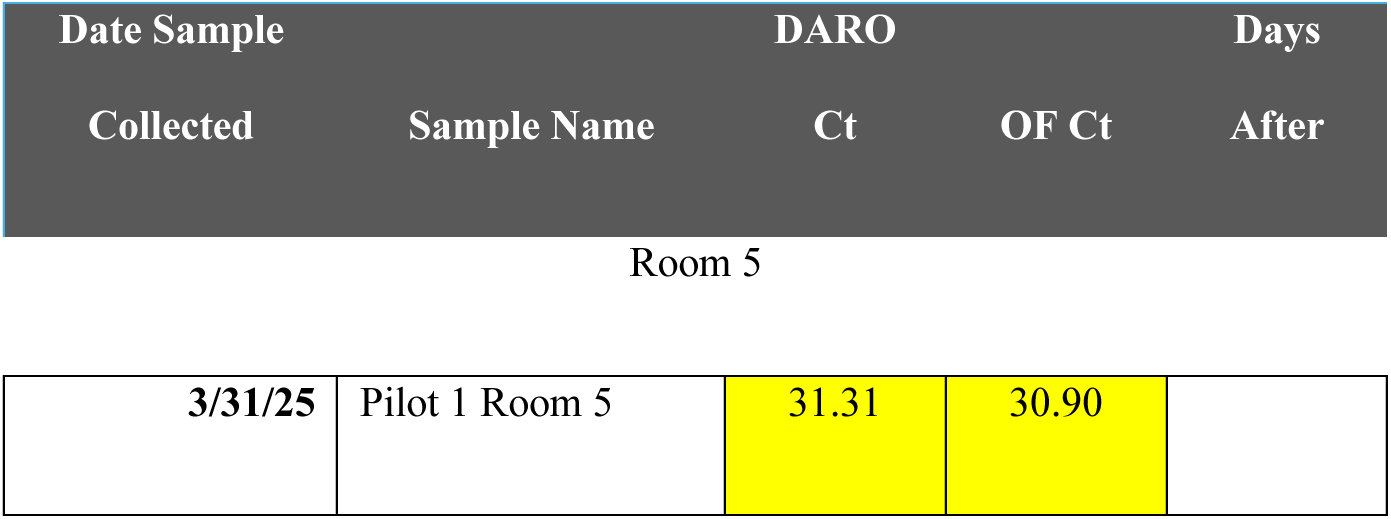
Room 5 within Pilot Study 1 was removed due to both oral fluid and DARO Systems testing identifying PRRSV on day 1 of the study.

**Table 4.**
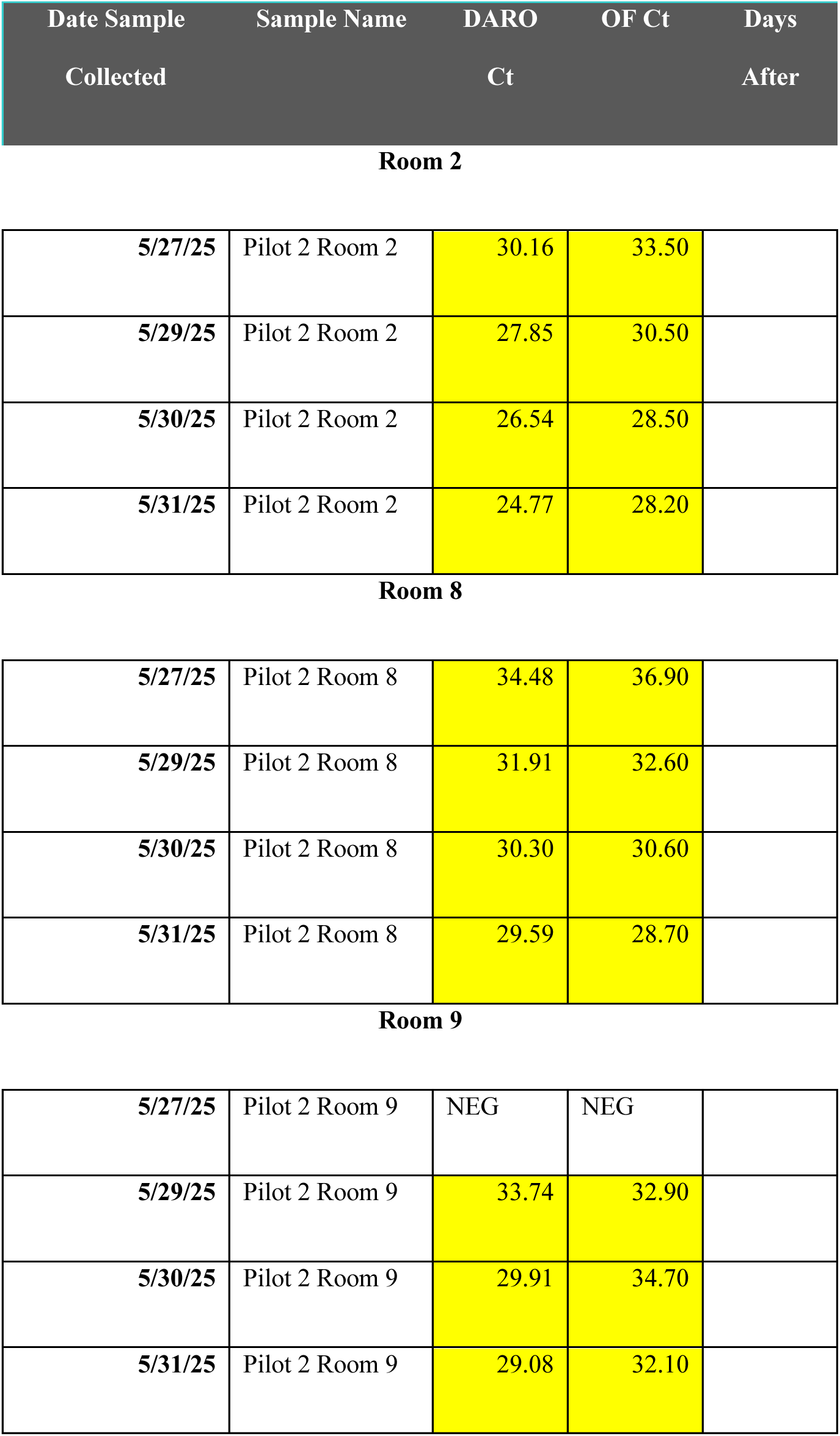
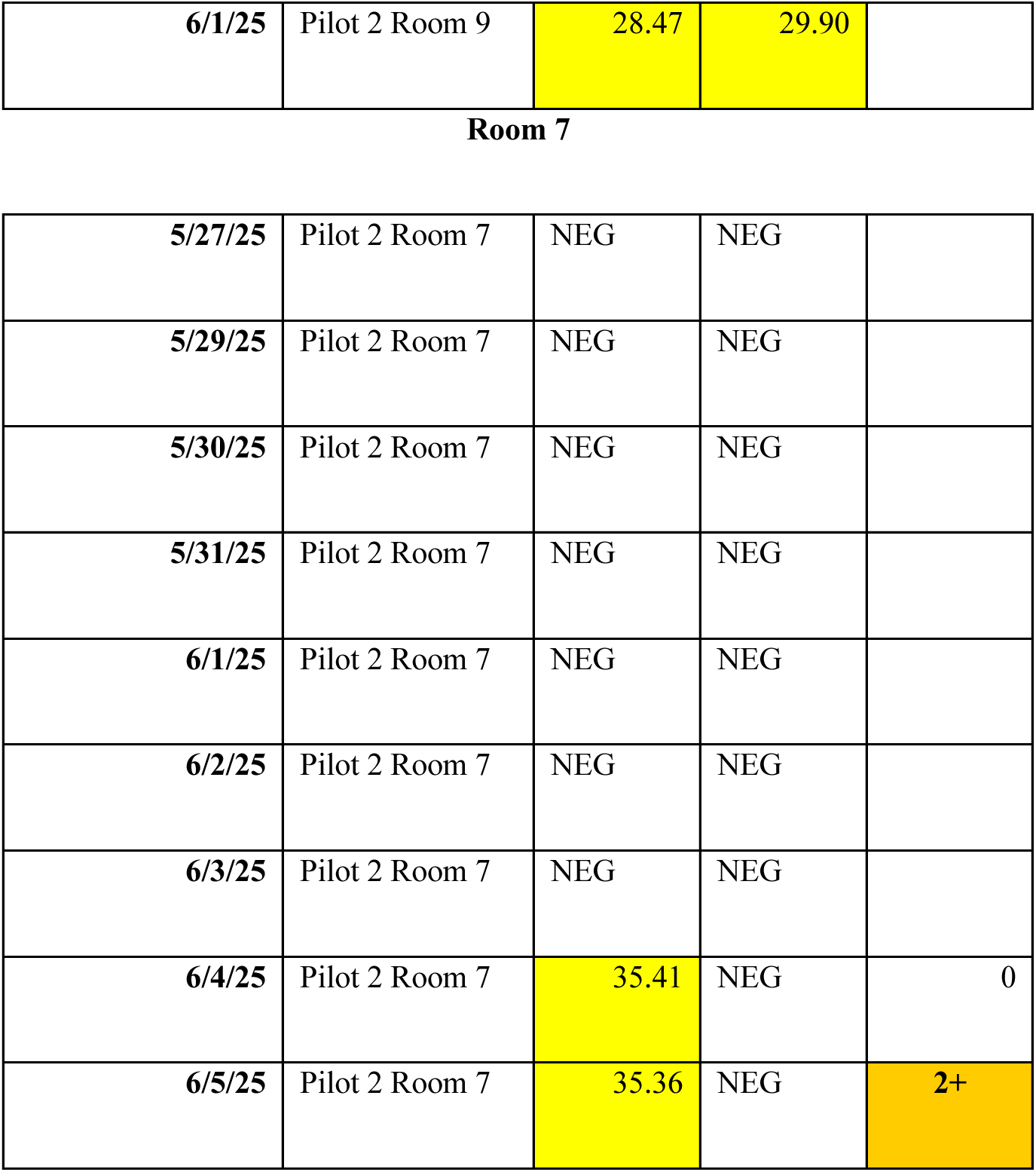
Room 2, Room 8, and Room 9 were removed from Pilot Study 2 analysis due to both oral fluid and DARO Systems testing identifying PRRSV on day 1 of the study. Additionally, Room 7 was removed from the analysis of the study as Oral fluids did not identify PRRSV presence.

**Room 2**
|  |  |  |  |
| --- | --- | --- | --- |
| 5/27/25 | Pilot 2 Room 2 | 30.16 | 33.50 |
| 5/29/25 | Pilot 2 Room 2 | 27.85 | 30.50 |
| 5/30/25 | Pilot 2 Room 2 | 26.54 | 28.50 |
| 5/31/25 | Pilot 2 Room 2 | 24.77 | 28.20 |
| **Room 8** |  |  |  |
| 5/27/25 | Pilot 2 Room 8 | 34.48 | 36.90 |
| 5/29/25 | Pilot 2 Room 8 | 31.91 | 32.60 |
| 5/30/25 | Pilot 2 Room 8 | 30.30 | 30.60 |
| 5/31/25 | Pilot 2 Room 8 | 29.59 | 28.70 |
| **Room 9** |  |  |  |
| 5/27/25 | Pilot 2 Room 9 | NEG | NEG |
| 5/29/25 | Pilot 2 Room 9 | 33.74 | 32.90 |
| 5/30/25 | Pilot 2 Room 9 | 29.91 | 34.70 |
| 5/31/25 | Pilot 2 Room 9 | 29.08 | 32.10 |
| **6/1/25** | Pilot 2 Room 9 | 28.47 | 29.90 |

**Room 7**
|  |  |  |  |  |
| --- | --- | --- | --- | --- |
| **5/27/25** | Pilot 2 Room 7 | NEG | NEG |  |
| **5/29/25** | Pilot 2 Room 7 | NEG | NEG |  |
| **5/30/25** | Pilot 2 Room 7 | NEG | NEG |  |
| **5/31/25** | Pilot 2 Room 7 | NEG | NEG |  |
| **6/1/25** | Pilot 2 Room 7 | NEG | NEG |  |
| **6/2/25** | Pilot 2 Room 7 | NEG | NEG |  |
| **6/3/25** | Pilot 2 Room 7 | NEG | NEG |  |
| **6/4/25** | Pilot 2 Room 7 | 35.41 | NEG | 0 |
| **6/5/25** | Pilot 2 Room 7 | 35.36 | NEG | 2+ |

All rooms throughout the study were designated a number rather than official name (i.e. Room 1, Room 2).

### Traditional surveillance collection and processing

Oral fluids were collected from designated pens using the swine producer’s protocol. A three-stranded ¾” diameter cotton rope was used for the oral fluids (OF) collection. The rope was unwound into three individual strands, and six of those strands were placed spatially throughout the room, distributed across 12 pens. Ropes were hung on gates at pig-height and pigs were allowed to chew for 30 minutes. The six strands were wrung out into a single 1-gallon plastic bag, manually homogenized, and 40 mL of the aggregated sample was poured into a 50 mL plastic tube. One tube was submitted per room sampled. Samples were stored at 4°C upon collection and immediately submitted for testing. Each tube was sent to Iowa State University Veterinary Diagnostic Lab (ISU VDL) by the producer and tested for PRRSV by reverse-transcriptase polymerase chain reaction (RT-PCR). Results and cycle threshold (Ct) values were reported to the producers and used for study evaluation. Samples were considered positive or negative by the lab-reported cutoff at a Ct value of ≤ 37.

### Novel surveillance collection and processing

Samples were collected using patent pending DARO Systems methodology (Pending Patent #19/256,939). Environmental fecal samples were collected across all pens in each room and pooled on a per room basis. Samples were placed on ice and stored at 4°C prior to testing. Collected samples were tested for the presence of PRRSV using patent pending DARO Systems sample processing technique (Pending Patent #19/256,939). Samples were prepped for nucleic acid extraction according to DARO Systems proprietary methodology to enrich for viral particles. Post preparation, samples were extracted using the MagMAX Wastewater Ultra Nucleic Acid Kit (Applied Biosystems, Foster City, CA, USA) according to manufacturer protocol. Post lysis of the sample, an internal positive control, Xeno (VetMAX Xeno internal positive control, Applied Biosystems, Foster City, CA, USA), was added to each sample at a concentration of 20,000 copies per reaction. Extracted samples were prepped for RT-qPCR using the VetMAX PRRSV 3.0 Reagent Kit (Applied Biosystems, Foster City, CA, USA) following the manufacturers protocol. Each RT-qPCR run included an extraction and template negative control as well as an extraction and template positive control. Runs were analysed using thresholds of 10,000 baseline of the target and internal Xeno control. Samples were considered positive with a Ct value of ≤ 39.

### Statistical Analysis

Statistical analyses were performed using the R package “survival” using the function “survfit” within R Studio version 2025.05.1+513 to evaluate statistical differences between DARO Systems and oral fluid sampling using a Kaplan-Meier Survival Analysis.^13, 14^ Additionally to evaluate proportional differences between DARO Systems and oral fluid surveillance testing “stats” with the function “mcnemar.test” within R Studio was used. Days to positive PRRSV result was used for the analyses. Graphs were generated using “ggplot2” within R Studio.^15^ All significance was determined at *P* < .05.

## Results

### A novel surveillance system shows rapid detection of PRRSV

Daily samples collected using the DARO Systems and OF Method were tested for the presence of PRRSV in each sampling room. Daily sampling started once either surveillance sampling approaches identified a positive sample and lasted until both surveillance sampling approaches tested positive for PRRSV. Both sites for Pilot 1 and Pilot 2 did have a natural PRRSV exposure and infection with the same 1-4-4 L1C.5.33 variant. Positive detection comparison identified DARO Systems detecting PRRSV on average 3.91 days ahead of Gold Standard OF (Figure 1 and Figure 2) throughout the duration of both pilot studies. Specifically, during Pilot 1, DARO Method detected PRRSV 4.14 days ahead of OF detection of PRRSV (Table 1, Figure 3). Similarly, PRRSV was detected 3.75 days earlier using DARO Systems methodology compared to OF sampling during Pilot 2 (Table 2, Figure 4). During Pilot 1, out of the eight rooms, one room tested positive for both OF and DARO Systems testing on day one and was removed from the study (Table 1). Similarly, during Pilot 2, three out of the eight rooms were omitted due positive results for both OF and DARO Systems testing on day one of the study and were removed from analysis (Table 2). One additional room (Room 7) from Pilot 2 was removed due to no positive identification of PRRSV via OF testing before shipment occurred.

**Figure 1.**
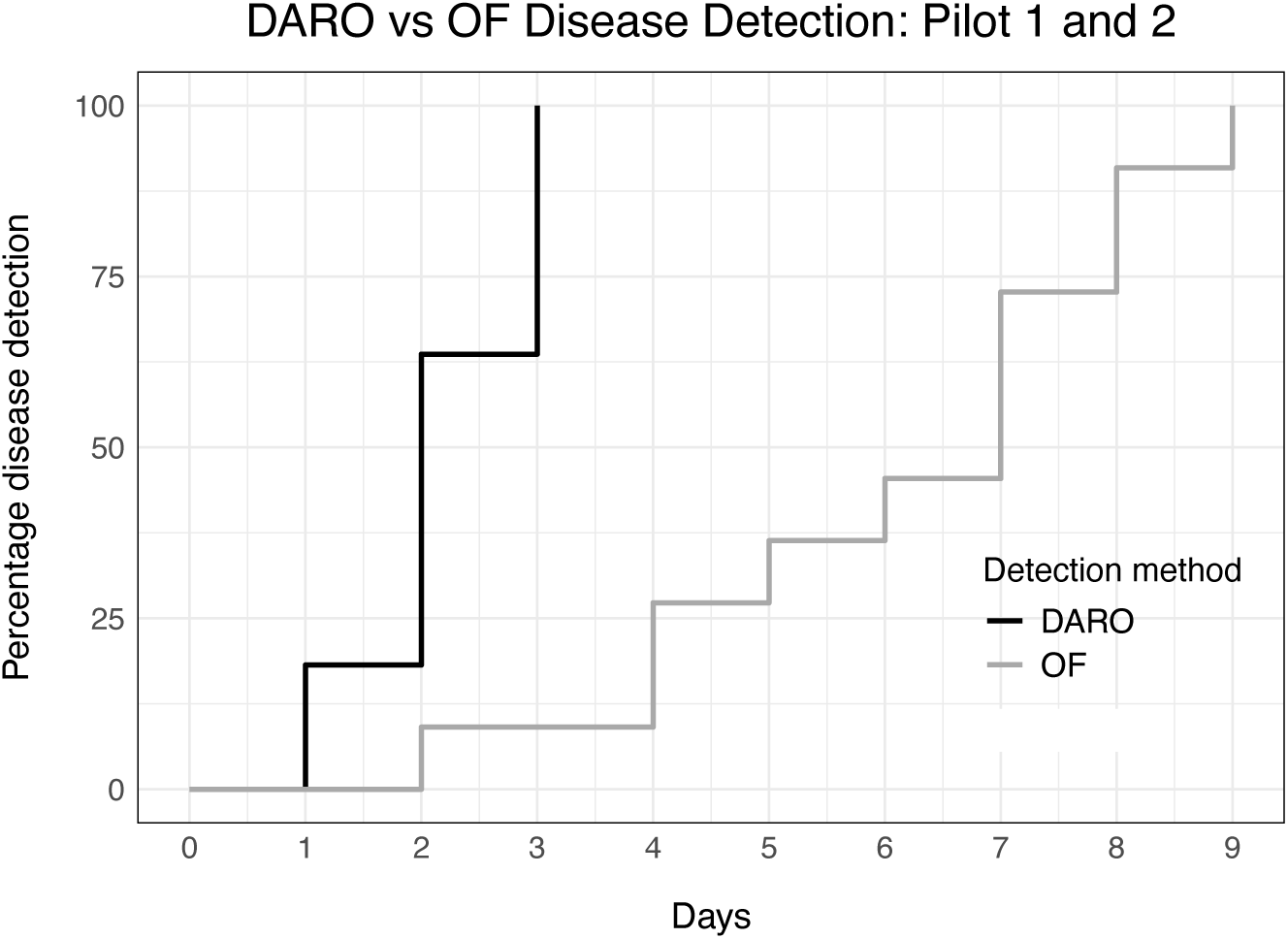
A Kaplan-Meier Survival Analysis was used to determine days until disease detection using both Pilot datasets (Pilot 1 and Pilot 2) where DARO Systems reached 100% by day 3 and oral fluids reached 100% by day 9. Statistical analysis showed a significant difference in days to a positive test between the two methodologies (*P* < .001).

**Figure 2.**
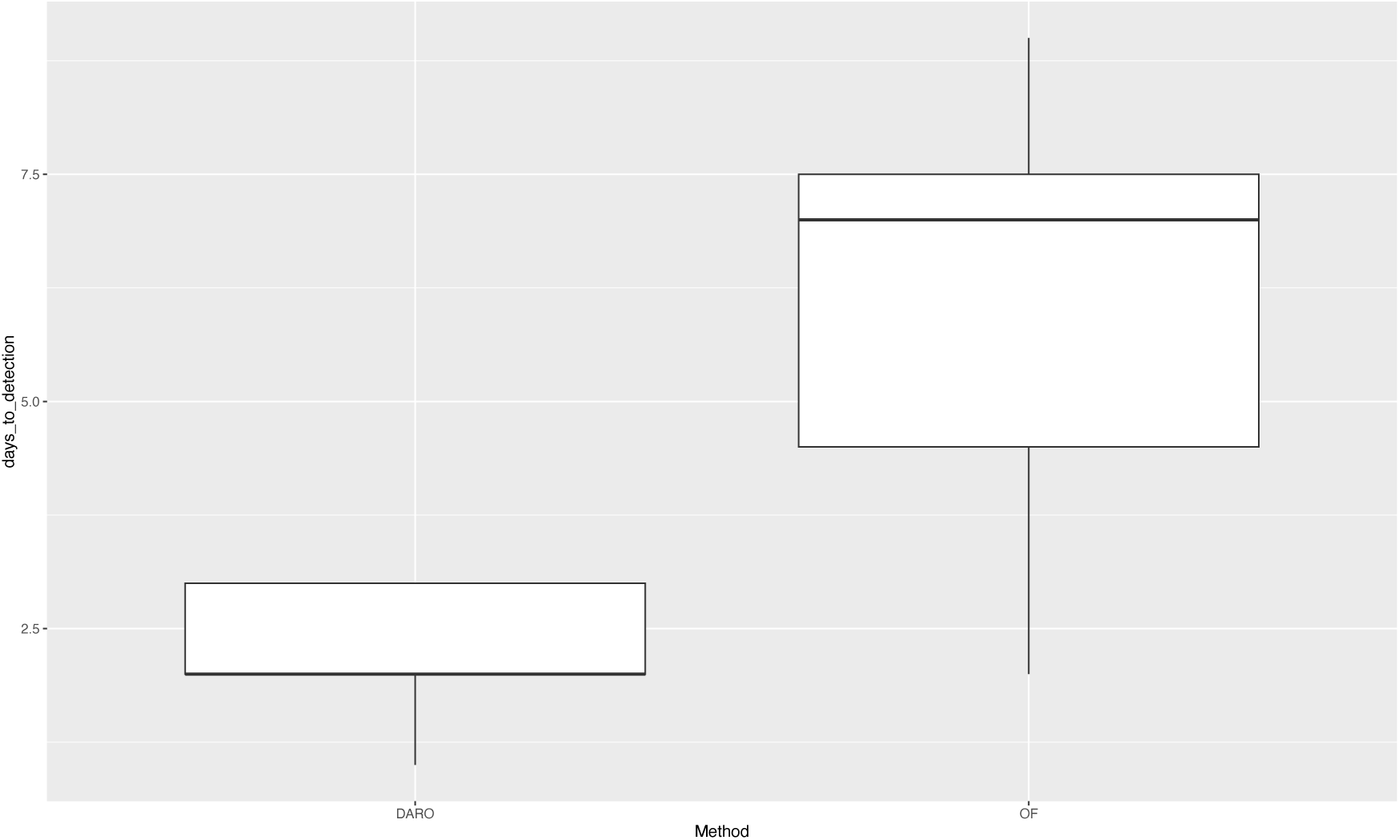
Box and Whisker analysis using both Pilot Data sets (Pilot 1 and Pilot 2) demonstrated DARO Systems detection of a PRRSV positive detection compared to oral fluid testing.

**Figure 3.**
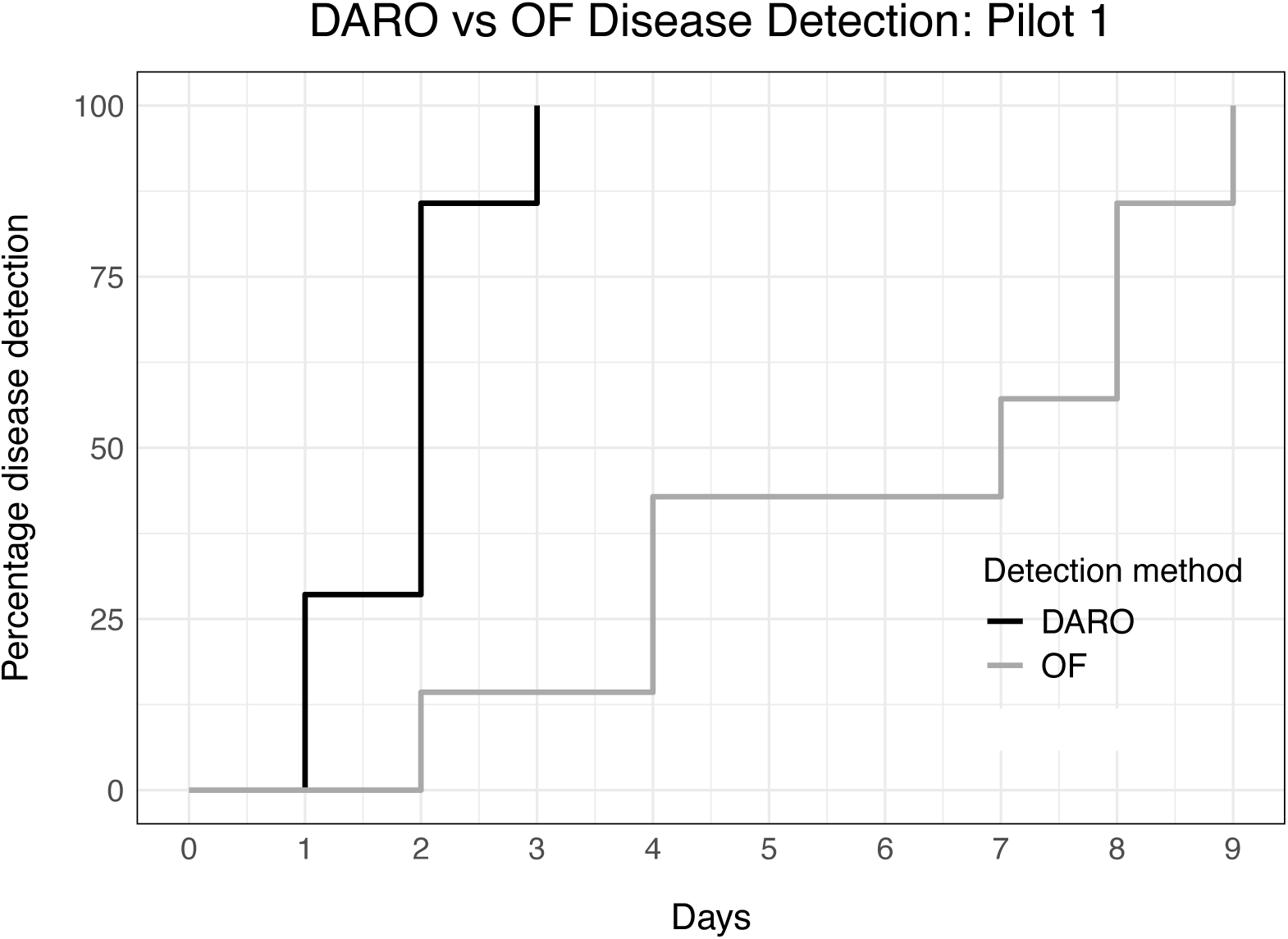
Pilot 1 data detecting PRRSV positive at 100% detection at day 3 of DARO Systems sampling compared to oral fluid 100% identification at day 9 using a Kaplan-Meier Survival Analysis. A significant difference was found at *P* = .001.

**Figure 4.**
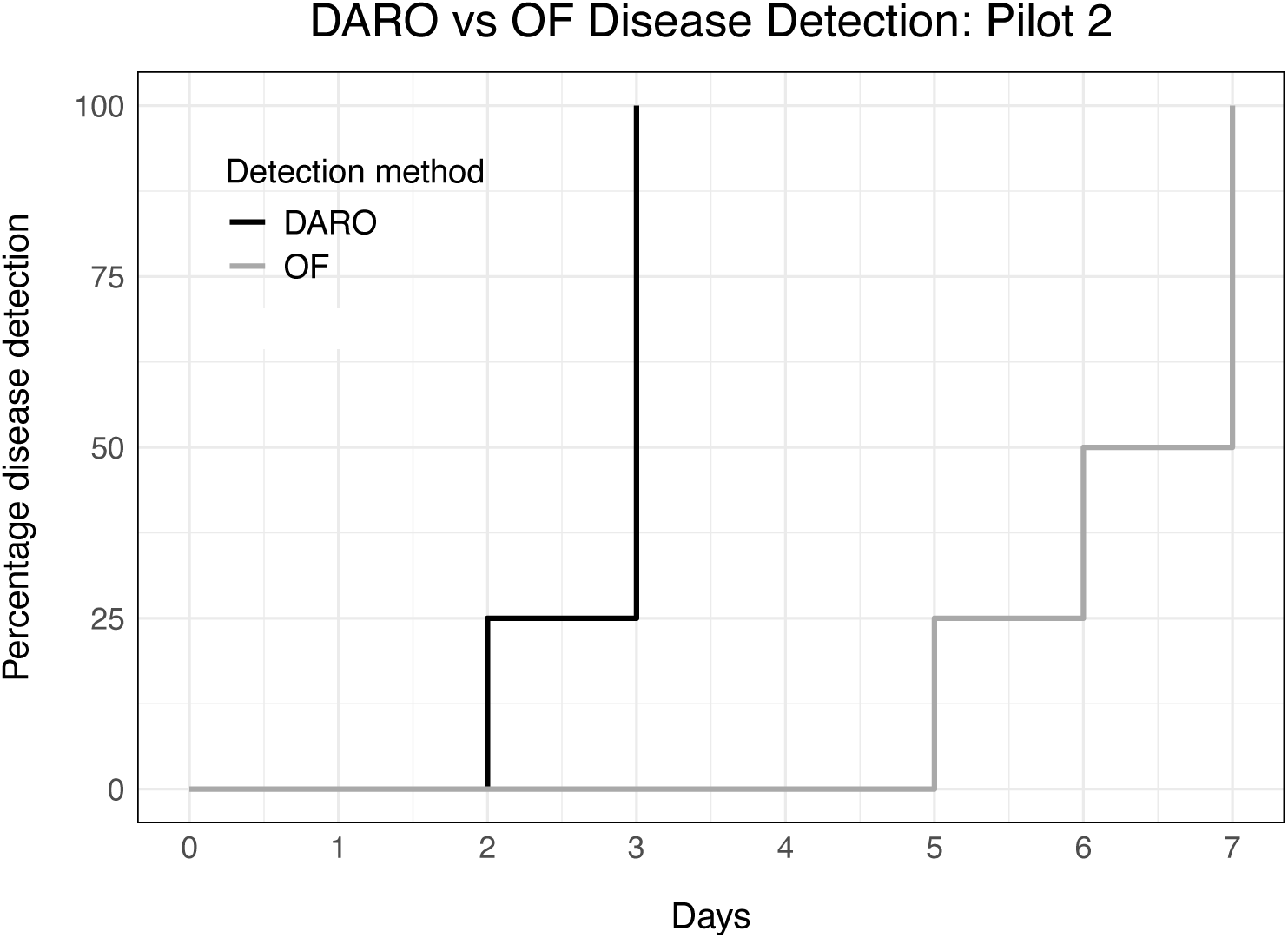
Pilot 2 data detecting PRRSV positive at 100% detection at day 3 of DARO Systems sampling compared to oral fluid 100% identification at day 7 using a Kaplan-Meier Survival Analysis. A significant difference was found at *P* = .01.

### Early-stage detection statistically differs

Statistical analyses further supported the observed differences between the two surveillance methodologies in time to a positive detection. A Kaplan–Meier survival analysis showed a significant difference in days to detection between the two surveillance methods in Pilot 1 (*P* = .001) and Pilot 2 (*P* = .01;). Across both data sets, days to detection differed significantly between the oral fluid and DARO Systems (*P* < .001; Figure 1). A Kaplan-Meier Survival Analysis showed the DARO Systems achieved 100% disease detection by day 3, whereas the oral fluid method reached 100% detection by day 9 (Figure 1). Consistent early disease detection trends were observed within each pilot where DARO Systems identified PRRSV prior to oral fluids. Specifically, in Pilot 1, DARO achieved 100% detection at day 3 compared to day 9 for oral fluids, and in Pilot 2, 100% detection was reached by day 3 for DARO and day 7 for oral fluids. A McNemars test further supports the observations seen between DARO Systems and oral fluids (*P* = .009). The 2X2 contingency table (Table 5) showed that out of 68 paired samples were 44 concordant while 24 were discordant (23 cases where DARO Systems was positive and oral fluids were negative; 1 case where oral fluid was positive and DARO Systems was negative).

**Table 5.** A McNemars test was used to investigate performance between the two surveillance collection methods.

|  | **Oral fluid +** | **Oral fluid -** |
| --- | --- | --- |
| **DARO Systems +** | 11 | 23 |
| **DARO Systems -** | 1 | 33 |

## Discussion

Utilizing two pilot studies, we compared a novel surveillance testing method with the DARO System to the industry-relevant standard, oral fluids, for the identification of PRRSV. Results showed earlier detection rate of PRRSV for DARO Systems compared to Gold Standard, oral fluids by an average of 3.91 days between the two pilot studies. Statistically significant difference was evaluated between the two sampling methodologies during Pilot 1 and Pilot 2 studies. Additionally, a McNemars test further demonstrates the significant difference between the two surveillance testing methods, suggesting a performance advantage to DARO Systems. Days of earlier detection rate for PRRSV presence is crucial for producers to make decisions on infection mitigation plans. The DARO Systems clearly exemplifies an opportunity for an extended window of time to provide producers to make these decisions. Specifically, these earlier detection times can allow for producers to make decisions for proactive treatment for potential secondary infections, herd closure for recovery, or in extreme cases, herd depopulation decisions. On a broader aspect, this novel collection and sampling method allows for producers to capture a non-invasive whole herd population test. Similarly, this surveillance method has potential to be implemented across multiple pathogen assays as well as multiple protein livestock species.

The use of feces for routine testing has shown great variability which in turn has led producers to utilize other specimens for routine testing. Excretion routes for PRRSV within feces shedding has been shown to be inconsistent.^16^ Studies show swine feces are composed of complex inhibitors that could be the causation of the inconsistency we see with using feces as a marker for pathogen shedding, such as PRRSV. Specifically, feces are known to be composed of but not limited to complex polysaccharides, bile salts, metabolic compounds, proteases, and DNases.^17^ The sample enrichment methodology developed within the DARO Systems (Patent Pending #19/256,939) effectively overcomes the challenge posed by fecal inhibitors, enabling accurate detection of excreted PRRSV. This study demonstrated the use of a whole-herd fecal collection method that surmounted the inconsistencies seen in fecal surveillance testing, even out-performing the industry standard oral fluids. Further the DARO Systems allows for a less invasive approach in sample collection (fecal collection vs blood), allowing for employees at any level to be able to obtain DARO Systems collected samples. This allows for additional producer savings in sample test cost as a veterinarian is not required to collect the sample.

### Limitations

While the DARO Systems shows great promise for its application in surveillance testing, there is a need for understanding limit of detection within this methodology. Additional sample size and location sites should be investigated for determination of days ahead for positive disease detection. Future directions include but are not limited to pooling studies to identify lower limit of detection, increased location site side by side testing, and increased user end awareness of accurate DARO Systems collection process will ensure viable samples for accurate results.

A potential limitation of the current study is the financial and intellectual property interest held by authors BMB, JLD, ACN, and KKB in DARO, Inc, the developer of the DARO Systems. This interest includes both equity in the company and status as inventors on a pending patent related to the technology. The authors affirm specific procedural safeguards were implemented to maintain objectivity. Data collections were performed by pre-defined protocols. Analysis was executed by Dr. John Nason who holds no financial interest in DARO, Inc. Additionally, the authors with the disclosed financial and intellectual property interests were not privy to the conventional side-by-side OF results until after the primary results from DARO Systems were finalized and reported. We believe these steps significantly mitigated the risk of conscious or unconscious bias influencing the data interpretation or reporting.

### Conclusions

Accurate and reliable early surveillance testing is needed to inform prompt treatment and mitigation strategies for PRRSV. Most frequent surveillance measures in growing pigs consist of aggregate population testing using oral fluid collections or to a lesser extent blood collection from individual swine^1^. However, blood collection is invasive and costly while oral fluid testing can lead to inaccurate results depending on animal behavior. This study comprised of two pilot studies to determine the efficacy of the DARO Systems compared to OF. Earlier PRRSV detection was consistently demonstrated using the DARO Systems over traditional OF sampling. As such, this novel surveillance system can detect PRRSV on average 3.91 days earlier than oral fluid testing.

Here, DARO Systems surveillance testing demonstrated an unbiased, non-invasive, whole-herd surveillance testing system that accurately and rapidly detects PRRSV earlier than traditional surveillance methodologies. On a broader impact, this system can be applied to other swine pathogens or additional livestock surveillance practices to ensure global food security.

### Implications

Under the conditions of this study:

- DARO Systems enables a more robust population based PRRSV sampling.
- Population based fecal surveillance identifies of respiratory viruses.
- DARO Systems gives an earlier window to act before disease spread.

## Acknowledgments

We would like to thank Dr. John Nason for his assistance in the statistical analysis.

## Disclaimer

Scientific manuscripts published in the *Journal of Swine Health and Production* are peer reviewed. However, information on medications, feed, and management techniques may be specific to the research or commercial situation presented in the manuscript. It is the responsibility of the reader to use information responsibly and in accordance with the rules and regulations governing research or the practice of veterinary medicine in their country or region.

